# iTOP: Inferring the Topology of Omics Data

**DOI:** 10.1101/293993

**Authors:** Nanne Aben, Johan A. Westerhuis, Yipeng Song, Henk A.L. Kiers, Magali Michaut, Age K. Smilde, Lodewyk F.A. Wessels

## Abstract

**Motivation:** In biology, we are often faced with multiple datasets recorded on the same set of objects, such as multi-omics and phenotypic data of the same tumors. These datasets are typically not independent from each other. For example, methylation may influence gene expression, which may, in turn, influence drug response. Such relationships can strongly affect analyses performed on the data, as we have previously shown for the identification of biomarkers of drug response. Therefore, it is important to be able to chart the relationships between datasets.

**Results:** We present iTOP, a methodology to infera topology of relationships between datasets. We base this methodology on the RV coefficient, a measure of matrix correlation, which can be used to determine how much information is shared between two datasets. We extended the RV coefficient for partial matrix correlations, which allows the use of graph reconstruction algorithms, such as the PC algorithm, to infer the topologies. In addition, since multi-omics data often contain binary data (e.g. mutations), we also extended the RV coefficient for binary data. Applying iTOP to pharmacogenomics data, we found that gene expression acts as a mediator between most other datasets and drug response: only proteomics clearly shares information with drug response that is not present in gene expression. Based on this result, we used TANDEM, a method for drug response prediction, to identify which variables predictive of drug response were distinct to either gene expression or proteomics.

**Availability:** An implementation of our methodology is available in the R package iTOP on CRAN. Additionally, an R Markdown document with code to reproduce all figures is provided as Supplementary Material.

**Supplementary information:** Supplementary data are available at *Bioinformatics* online.

## 1 Introduction

Rapid developments in high throughput measurement techniques together with rapid reduction in profiling costs have, for many biological problems, endowed us with multiple molecular datasets recorded on the same set of objects. For example, pharmacogenomics data contain, in addition to cancer type and drug response, various omics datasets (mutation, copy number aberration (CNA), methylation, gene expression and proteomics) recorded on the same set of tumor cell lines (4; 6). While this provides an unprecedented view on the underlying biological problem, it also comes with some unique challenges. Specifically, the recorded datasets are not independent of each other, but are characterized by specific relationships. For example, copy number alterations and methylation changes may influence gene expression, which may, in turn, influence drug response. As we have demonstrated earlier (1), these relationships can have profound effects on further integrative analyses, especially biomarker discovery. It is therefore imperative to obtain a full quantitative characterization of these relationships, such as the illustrative topology of relationships between datasets depicted in Figure 1A.

**Fig. 1.**
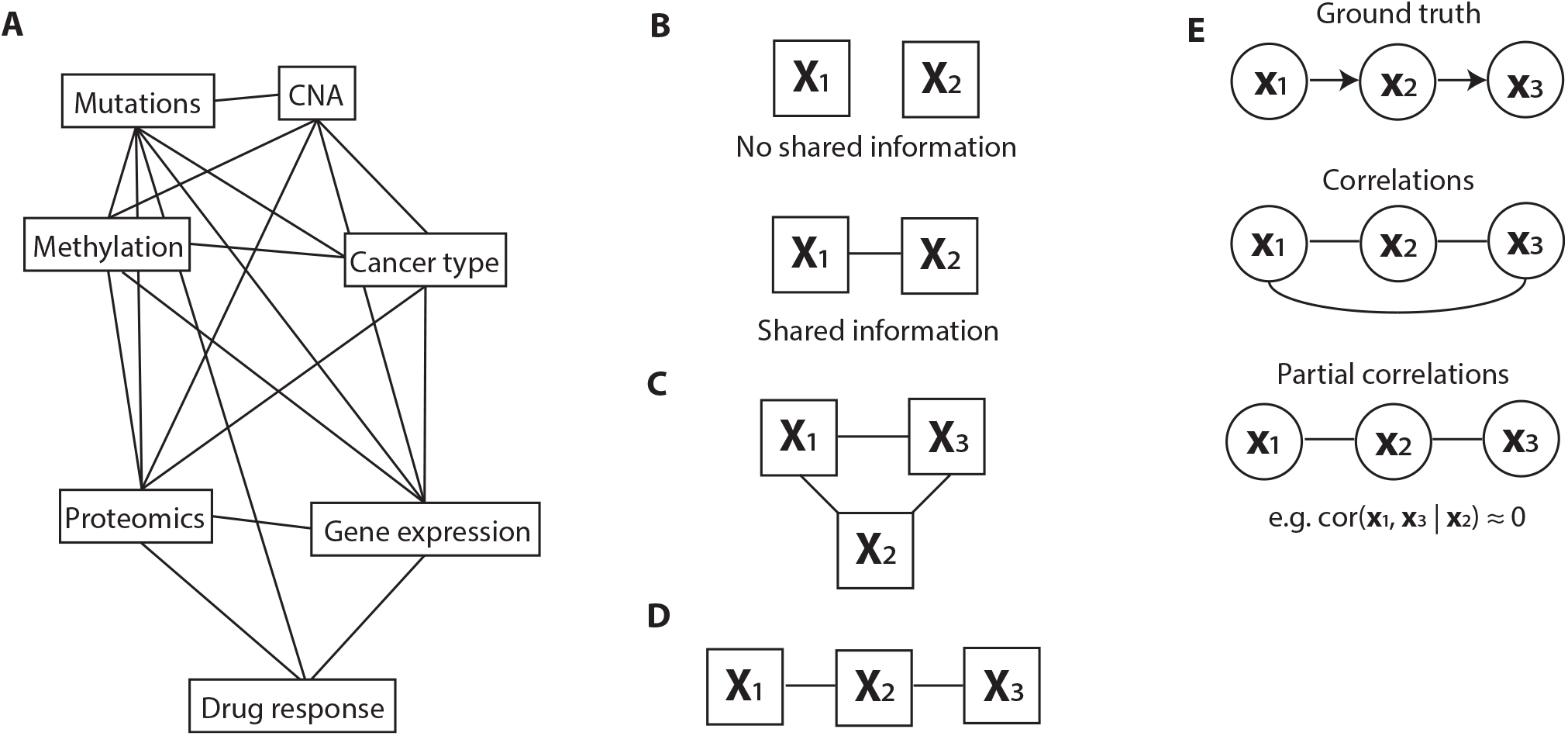
High-level overview of this work. (A) The goal of this work is to infer a topology of relationships between pharmacogenomics datasets (an example topology is illustrated here). (B) When two datasets share information (i.e. when their RV coefficient is non-zero), we will indicate them as connected in a topology. (C) A topology of three datasets that all share information. We will convert this topology to the one depicted in (D) if the shared information between X_1_ and X_3_ is fully contained in X_2_. (E) To create these topologies we will draw on methods for inferring a topology between single variables using partial correlations. Top: the original causality graph. Middle: the topology as inferred using correlations. Bottom: the inferred topology using partial correlations.

Here we set out to characterize the relationships between datasets in terms of the amount of information that is shared between a pair of datasets, and, more importantly, how this shared information manifests itself in the relationship of a pair of datasets to a third dataset. For example, suppose we have two datasets, **X**_1_ and **X**_2_. Suppose we can characterize the amount of shared information between **X**_1_ and **X**_2_ by a number between 0 and 1, with 0 being no shared information and 1 representing maximal overlap in information (Figure 1B). This characterization of pairwise relationships can be informative as such, as it can reveal whether, for example, there is any shared information between gene expression and mutation data. If we now introduce a third dataset, **X**_3_, we can also quantify the amount of information shared between **X**_1_ and **X**_3_ and **X**_2_ and **X**_3_. Assuming that these relationships are non-zero, we obtain the graph in Figure 1C. Now it becomes particularly interesting to know whether the shared information between **X**_1_ and **X**_3_ depends on **X**_2_. Specifically, is the shared information between **X**_1_ and **X**_3_ contained in the information in **X**_2_?In other words, does **X**_2_ mediate the effect of **X**_1_ on **X**_3_? When these questions can be answered for all datasets at hand, it reveals the minimal graph that represents the conditional relationships between all datasets. As the number of datasets grows, such a graph not only gives a very concise overview of the relationships, but it is also an important guide in structuring the analyses aimed at finding biomarkers of a given phenotype. More specifically, suppose that **X**_1_, **X**_2_ and **X**_3_ represent mutation, gene expression and drug response data for a cell line panel, and that our goal is to extract molecular biomarkers of drug response. Assume that, from our analyses, it emerged that all the information shared between mutation (**X**_1_) and drug response (**X**_3_) is contained in the gene expression data (**X**_2_) (Figure 1D). This implies that we only need to employ gene expression data to find biomarkers of drug response.

To infer dataset topologies, we draw upon the approaches employed to infer topologies between single variables (instead of matrices). Specifically, for our earlier example, we can employ partial correlation, e.g. *cor*(**x**_1_, **x**_3_|**x**_2_), to quantify the amount of information that is shared between two variables (**x**_1_ and **x**_3_) that is not present in the other variable (**x**_2_). If the effect of **x**_1_ on **x**_3_ is (almost fully) mediated through x_2_, it follows that *cor*(**x**_1_, **x**_3_|**x**_2_) ≈ 0, which implies that we can remove the direct link between x_1_ and x_3_ (Figure 1E). Graph reconstruction algorithms, such as the PC algorithm (12; 2), use this property to infer the topology between multiple variables.

Here, we propose iTOP, a methodology for inferring topologies between datasets. As with topology inference for single variables, this methodology consists of two components: 1) a measure of (conditional) similarity between datasets and 2) the PC algorithm that employs the (conditional) similarity measure to perform structure learning, i.e. to infer the topology. As similarity measure we employ the RV coefficient (9), a measure of matrix correlation. The basic idea of the RV coefficient is that datasets are correlated when they have a similar configuration (e.g. similar clustering) of the objects. We extend the RV coefficient to be applicable to binary data by using Jaccard similarity to determine the configuration of objects. This allows us to measure the shared information between any of the molecular datasets, including intrinsically binary datasets such as mutation data. In addition, to measure conditional matrix similarity, we extend the RV coefficient for partial matrix correlations. This allows us to quantify the amount of information that is shared between two *datasets* (matrices), but not present in the other dataset, analogous to single variables.

We employ iTOP, i.e. partial matrix correlation in conjunction with the PC algorithm, to infer a topology of relationships between datasets. First, we will demonstrate the RV coefficient with both extensions (i.e. for partial matrix correlations and for binary data) on artificial data. Subsequently, we will use this to infer the topology of relationships between the pharmacogenomics datasets. We show that gene expression acts as a mediator between most other datasets and the drug response, and that only proteomics clearly shares information with drug response that is not present in gene expression. Based on this result, we will employ TANDEM, a method for drug response prediction from multiple datasets (1), to identify markers predictive of drug response that are distinct for proteomics and gene expression.

## 2 Methods and Materials

### 2.1 Matrix correlation using the RV coefficient

For dataset *i,* consider **X**_*i*_ the *n* × *p*_*i*_ data matrix with objects in the rows and variables in the columns. Here, we assume **X**_*i*_ to be column-centered (of note, there is no need to scale the columns of **X**_*i*_). We define the corresponding *n* x *n* configuration matrix **S**_*i*_ as follows:

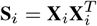

Now consider a second dataset *j,* whose data matrix **X**_*i*_ has the same objects on the same rows as **X**_*i*_, but has a different set of variables. Hence, **X**_*j*_ is of size *n* × *p*_*j*_. Analogous to **X**_*i*_, we will define a configuration matrix **S**_*j*_ for **X**_*i*_.

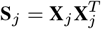

Using the configuration matrices **S**_*i*_ and **S**_*j*_, we can then determine the matrix correlation between these matrices using the RV coefficient:

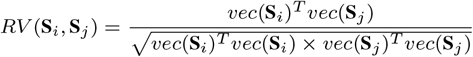

Where *vec*(**S**) is the *n*^2^ × 1 vector in which the columns of **S** are stacked on top of each other. When **X**_*i*_ and **X**_*j*_ are column-centered, then *mean*(*vec*(**S**_*i*_)) = 0 and *mean*(*vec*(**S**_*j*_)) = 0, which means we can interpret the above as a Pearson correlation coefficient.

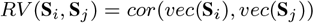

### 2.2 The modified RV coefficient

For data matrices **X** where the number of variables is much greater than the number of objects (i.e. *p* ≫ *n*), the RV coefficient is known to be biased upwards (10; 8). To account for this bias, we subtract the diagonal from the configuration matrix, as in the modified RV coefficient (10).

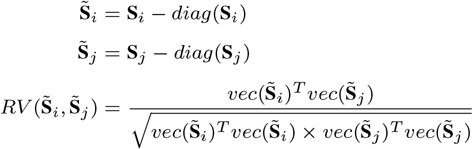

For a more complete discussion of the modified RV coefficient, as well as our rationale for not using the adjusted RV coefficient (8) instead, we refer to the Supplementary Material.

### 2.3 Partial matrix correlations

We extend the above matrix correlation formulation to partial matrix correlations. Consider a third dataset, the *n* × *p*_*k*_ matrix **X**_*k*_, that will be processed as above.

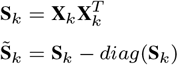

We can then compute the partial matrix correlation between dataset *i* and *j,* corrected for dataset *k,* as

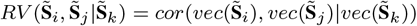

Of note, the concept of partial matrix correlations has been explored previously by Smouse et al. (1986) (11), who based their measure on the Mantel Test (7). For a discussion of the Mantel Test and why we prefer to base our measure of partial matrix correlation on the RV coefficient, we refer to the Supplementary Materials.

### 2.4 Statistical inference for partial matrix correlations

We provide two methods for statistical inference for partial matrix correlations: significance estimates and confidence intervals. We note that these cannot be determined analytically (e.g. using Fisher Transformation, which is commonly used to derive a p-value for Pearson correlations), as the entries in *vec*(**S**) are not i.i.d.: multiple entries in *vec*(**S**) correspond to the same object in **S**. Instead, we will discuss a permutation test for significance estimates and a bootstrapping procedure for calculating confidence intervals.

We used a permutation test to assess significance of a (partial) matrix correlation. In every permutation, the objects of every dataset were independently shuffled and the (partial) matrix correlation was computed on the shuffled data. Subsequently, the observed (partial) matrix correlation was compared to the permuted values, and the p-value was set to

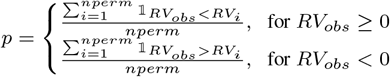

Where 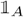 is the indicator function that equals 1 when *A* is true, *RV*_*obs*_ is the observed (partial) matrix correlation, *RV*_*i*_ the permutated (partial) matrix correlation from the *i*th permutation and *nperm* the number of permutations. Throughout the manuscript, we used *nperm* = 1000.

We used a percentile bootstrap procedure to calculate confidence intervals. In each bootstrap, objects were obtained by drawing complete cases randomly (with replacement) from the dataset, after which the (partial) matrix correlation was calculated as defined above. The 99% percentile interval of the obtained (partial) matrix correlations was then used as a confidence interval. Throughout the manuscript, we used 1000 bootstraps to determine a confidence interval.

We note that row-wise permutation of the data matrices (**X**[*ind*,], with *ind* the indices of the objects after permutation) is equivalent to permutation of both the rows and the columns of the configuration matrices (**S**[*ind*, *ind*]). Using this property, we decided to permute the configuration matrices, as this prevents having to calculate the configuration matrix in each permutation and hence greatly speeds up the calculations. A similar approach was used for bootstrapping.

### 2.5 Binary similarity measures

An advantage of converting the data matrices **X** to configuration matrices **S** is that it allows us to use different similarity measures for different data types. For example, for continuous data, we use:

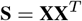

Note that each entry of **S** corresponds to an inner product between different objects in **X**, i.e.

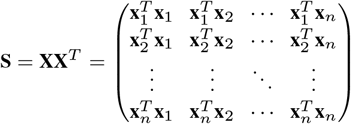

Where **x**_*i*_ is the *i*’th row in **X** and *n* is the number of rows in **X**.We will refer to this similarity measure as ‘inner product similarity’.

#### 2.5.1 Jaccard similarity

For binary data, we use Jaccard similarity. Jaccard similarity is defined as the ratio of the number of elements where these vectors have ones in common and the total number of positions where ones occur in any of these two vectors. Consider the following contingency table.

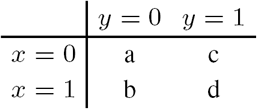

Where *a* is the number of elements where *x =* 0 and *y =* 0, *b* is the number of elements where *x =* 1 and *y =* 0, etc. The Jaccard Similarity can then be written as:

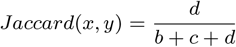

When all *x =* 0 and all *y =* 0, then *b = c = d =* 0, which would result in 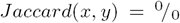. In these cases, we define the Jaccard similarity as *Jaccard*(*x*, *y*) = 0.

Note that the Jaccard similarity is based on the number of positive matches (*d*) and not at all on the number of negative matches (*a*). This is in line with our intuition of similarity in the binary data at hand (mutation, CNA and cancer type). For example, when two objects share the same mutations, we think this should contribute more to their similarity than the number of mutations that both objects lack.

We define configuration matrices using the Jaccard similarity in the following way:

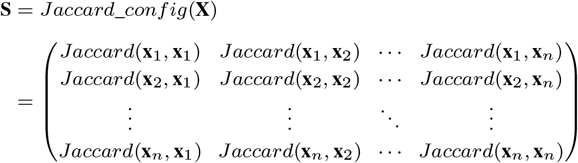

#### 2.5.2 Kernel centering

We used kernel centering to center the configuration matrix **S** rather than the underlying data matrix **X**. Essentially, kernel centering is double centering (i.e. column- and row-wise centering) of the configuration matrix **S** (or in other words: the kernel), which we will show to be equal to first column-centering the data matrix **X** and then computing **S** = **XX**^*T*^. Consider **X** the original data matrix and 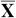 the column-centered data matrix. Likewise, consider S the original configuration matrix and 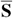 the centered configuration matrix. Finally, consider **m** the column-wise means of **X** and *n* the number of rows in **X**. We will first consider an example using inner products as a similarity measure.

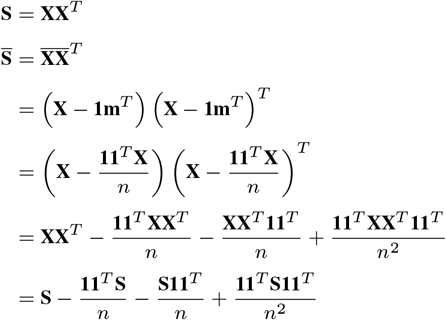

Interestingly, the final term expresses the kernel centered 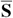 in terms of the non-centered **S**. This allows us to center configuration matrices that are not based on inner-product similarity, such as **S** = *Jaccard_config*(**X**). Column-centering **X** (the input space) makes no sense here, as the resulting matrix would not consist of 0s and 1s anymore and hence 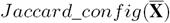 is not defined. However, we can use kernel centering here to center the so-called kernel space corresponding to **S**.

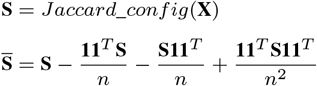

### 2.6 Pharmacogenomics data

The mutation, copy number aberration (CNA), methylation, cancer type, gene expression and drug response data were sourced from GDSC1000 (4), and the proteomics data were sourced from MCLP (6) (Table 1). For the mutation and CNA data, we used the reduced set of Cancer Functional Events (CFEs) (4), resulting in 300 and 425 binary variables respectively. For the methylation data, we used the CpG-island summarized data, resulting in 14,426 continuous variables. For the cancer type data, we used the classification into 30 TCGA cancer types or ‘OTHER’, resulting in 31 binary variables (4). For gene expression data, we used the gene level summarizeddata, resultingin17,419continuous variables. The proteomics data consist of 452 variables, of which 108 represent phospho-protein levels and the remaining 344 represent protein abundance levels. For the drug response data, we used the IC50-values (concentration at which half of the cells are killed) for all 265 drugs.

**Table 1.** Overview of the pharmacogenomics datasets used in this manuscript.

|  | Dimensionality | Source | Type | Missing values |
| --- | --- | --- | --- | --- |
| Mutation | 300 | GDSC1000 | Binary | No |
| CNA | 425 | GDSC1000 | Binary | No |
| Methylation | 14,429 | GDSC1000 | Continuous | No |
| Cancer type | 31 | GDSC1000 | Binary | No |
| Gene expression | 17,419 | GDSC1000 | Continuous | No |
| Proteomics | 452 | MCLP | Continuous | Yes |
| Drug response | 265 | GDSC1000 | Continuous | Yes |

Of the 282 cell lines that were profiled in both GDSC1000 and MCLP, 266 cell lines were characterized across all seven datasets. This number was further reduced due to missing values in the proteomics and drugs response data. For the proteomics data, after removing all variables with >30% missing values, we retained 186 variables. Subsequently, after removing all objects with >30% missing values, we retained 221 objects. We then intersected all datasets with these 221 objects and applied the same two steps to the drug response data, where we retained 206 objects and 217 variables. These 206 objects cover 27 of the 31 cancer types in the GDSC1000 data. The remaining missing values (1% for the proteomics and 5% for the drug response) were imputed using SVD imputation (13) as implemented in the R package bcv.

## 3 Results

### 3.1 The RV coefficient

To illustrate the RV coefficient, consider the following example. Figure 2A represents data matrix **X**_1_, a dataset with two variables and 100 objects, where the first 50 objects form the green cluster and the second 50 objects form the purple cluster. The second data matrix, **X**_2_ (Figure 2B), also consists of two variables and the same 100 objects with the same clustering as in **X**_1_. The third data matrix, **X**_3_ (Figure 3C), is again a dataset with two variables and the same objects as before, but now without any apparent clustering.

**Fig. 2.**
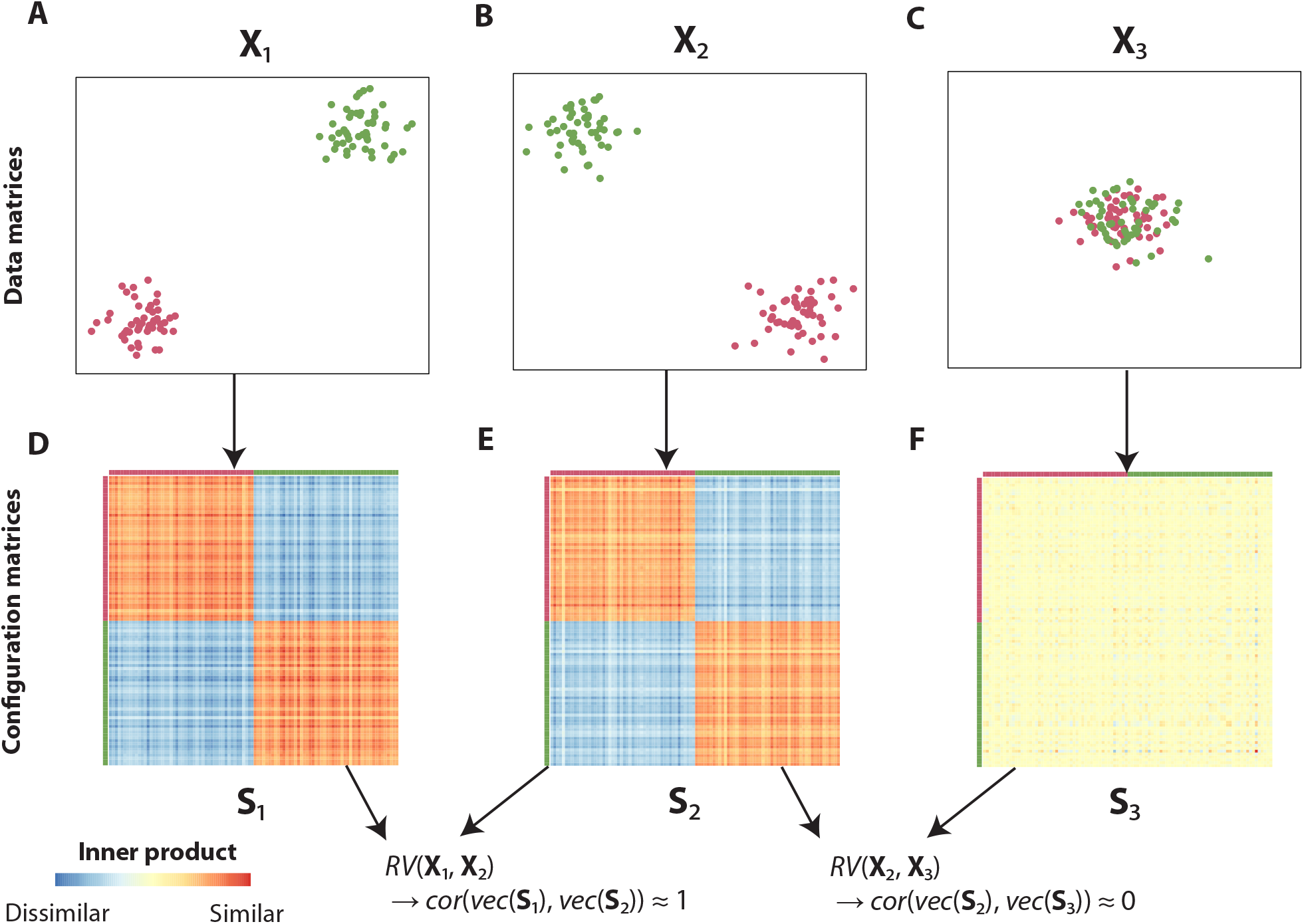
The RV coefficient explained using three simple example datasets. The data matrices X_1_, X_2_ and X_3_ (represented in A-C) are converted to configuration matrices S_1_, S_2_ and S_3_ respectively (D-F). Using the configuration matrices, it can be readily seen that *RV* (X_1_, X_2_) ≈ 1 and *RV* (X_1_, X_3_) ≈ 0.

When converting these data matrices to configuration matrices (similarity matrices), which indicate the configuration of the different objects with respect to each other, it can be readily observed that **X**_1_ and **X**_2_ contain the same information in terms of clustering (Figure 2D & E). Indeed, when computing the RV coefficient between **X**_1_ and **X**_2_ (by computing the Pearson correlation of the vectorized forms of the corresponding configuration matrices, see Methods and Materials), we obtain an RV coefficient close to one, indicating a strong relationship. Conversely, when computing the RV coefficient between **X**_2_ and **X**_3_, where the latter contains no clustering information, we see that the configuration matrices are very different and *RV*(**X**_2_, **X**_3_) ≈ 0 (Figure 2C & F).

### 3.2 Extending the RV coefficient for partial matrix correlations

We illustrate partial matrix correlations using the following example. Consider three datasets: **X**_1_, **X**_2_ and **X**_3_. Let **X**_1_ affect **X**_2_, and let **X**_2_ affect **X**_3_ (Figure 3). Observe that, consistent with the proposed causality, **X**_1_ is most similar to **X**_2_ (only the purple cluster in the bottom-left has been moved) and **X**_3_ is most similar to **X**_2_ (only the blue cluster in the bottom-right has been moved). This of course means that *RV*(**X**_1_, **X**_2_) and *RV*(**X**_2_, **X**_3_) will be non-zero. However, note that also *RV*(**X**_1_, **X**_3_) will be non-zero, as **X**_1_ and **X**_3_ do share information: the top three clusters have the same configuration in both datasets. Therefore, if we were to infer a topology based on the matrix correlations, we cannot rule out a direct link from **X**_1_ to **X**_3_.

Using the partial matrix correlation *RV*(**X**_1_, **X**_3_|**X**_2_), we can rule out a direct link from **X**_1_ to **X**_3_. As **X**_2_ has the same configuration in the top three clusters, correcting for **X**_2_ results in *RV*(**X**_1_, **X**_3_|**X**_2_) = 0.005 *≈* 0. Therefore, using partial matrix correlations, we can indeed reconstruct the original topology.

### 3.3 Extending the RV coefficient for binary data

The RV coefficient has been proposed for comparing data matrices containing continuous values. Specifically, in the original formulation of the RV coefficient, the configuration matrices are determined using the inner product between objects (Methods and Materials), which is tailored to comparing continuous values. To determine (partial) matrix correlations for datasets containing binary values, we propose to create the configuration matrices using Jaccard similarity, which determines similarity between binary variables (Methods and Materials). We assessed the performance of this approach using a simulation study.

First, to establish a reference, we performed a simulation study in which two continuous valued matrices were compared. In this simulation, the values in **X**_1_ and **X**_2_ were randomly drawn from *N(*10, 1) and *N*(0, 1) respectively, where *N*(*µ*, *σ*) represents a Gaussian distribution with mean *µ* and standard deviation *σ.* Subsequently, we defined a third matrix as **X**_3_ = (1 – α)**X**_1_ +α**X**_2_. We compared *RV*(**X**_1_, **X**_3_) for different values of *α,* and both with and without column-wise centering of the data matrices (Figure 4A). Regardless of centering, we found that *RV*(**X**_1_, **X**_3_) = 1 for *α =* 0 and *RV*(**X**_1_,**X**_3_) ≈ 0 for *α =* 1, as expected. For intermediate values of *α* however, we see big differences between the approach using centering and the one without centering. Without centering, *RV*(**X**_1_,**X**_3_) remains very close to 1 for values of α approaching 1, which is counterintuitive. With centering, *RV*(**X**_1_, **X**_3_) slowly decreases to 0 as *α* increases, which is according to expectation. These differences can be attributed to the fact that inner product distance is dependent on the relative position of the objects with respect to the origin: in the uncentered case, for *α* ≤ 0.9, the vectors representing the objects in **X**_1_ and **X**_3_ will be highly collinear, resulting in an RV coefficient close to one (Supplementary Figure 1). This experiment emphasizes the importance of centering the data prior to applying the RV coefficient.

**Fig. 3.**
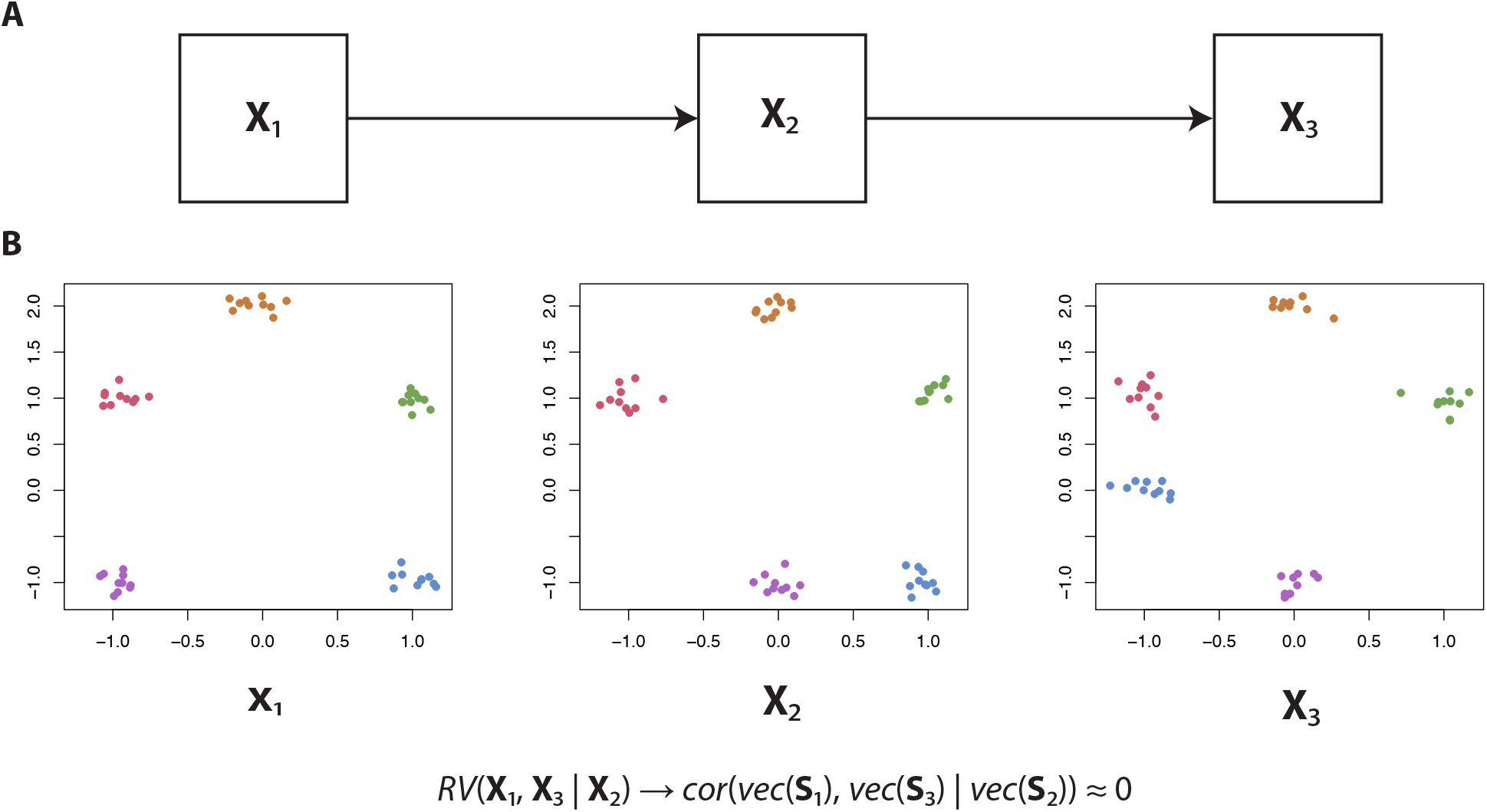
Illustration of the partial matrix correlation. (A) We will create artificial data such that X_1_ influences X_2_, which in turn influences X_3_. (B) Artificial data consistent with the abovementioned causality, resulting in *RV*(X_1_, X_3_ |X_2_) ≈ 0.

We then performed a simulation in which two binary valued matrices were compared. Values in **X**_1_ were randomly drawn from *Binom*(0.5) (Binomial distribution with *p =* 0.5). **X**_2_ was set equal to **X**_1_, but with *α* the fraction of binary values that were flipped. We varied *α* only up to 0.5, as this is the point at which the configuration of objects in **X**_1_ and **X**_2_ is maximally apart (at *α =* 1, **X**_1_ and **X**_2_ are simply inverted and, given that the RV coefficient is rotation independent, the resulting RV coefficient will be 1 again). Again *RV*(**X**_1_, **X**_2_) was compared for different values of *α* and both with and without centering (Figure 4B). As binary data cannot be column centered (it would not be binary anymore after centering), we instead used kernel centering to center the configuration matrix obtained using the Jaccard similarity (Methods and Materials). For *α =* 0, *RV*(**X**_1_, **X**_2_) = 1, both with and without centering, as the two matrices are exactly the same. However, for *α* in (0,50], *RV*(**X**_1_, **X**_2_) remained very close to 1 in the uncentered case, while it slowly decreased to 0 in the centered case. Hence, as at *α =* 0.5 the configuration of **X**_1_ and **X**_2_ is maximally apart, the centered case is preferable.

**Fig. 4.**
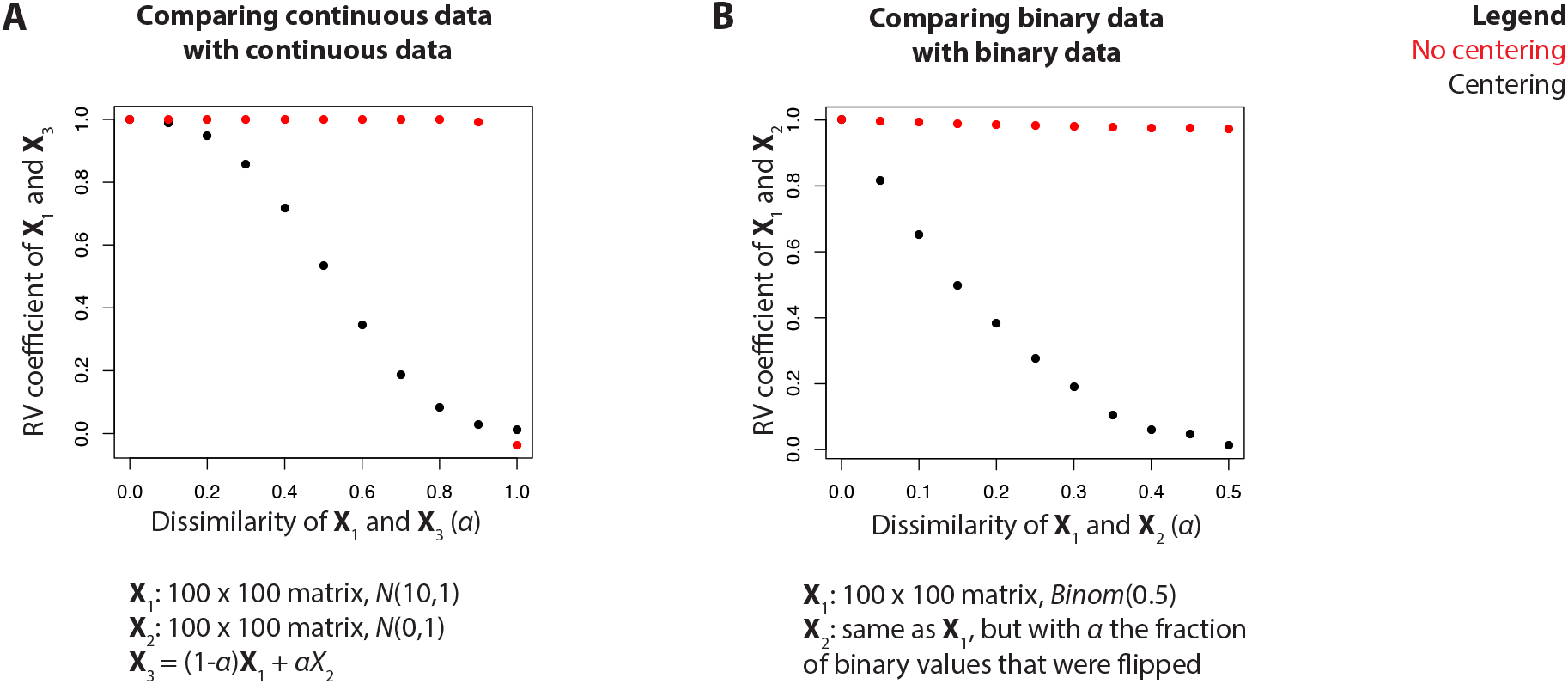
Artificial data experiment in which the RV coefficient (*y*-axis) is measured at different levels of similarity (*α, x*-axis), both with and without centering, for (A) two continuous datasets and (B) two binary datasets.

Using these simulation experiments, we have shown that the Jaccard similarity can be used to construct configuration matrices for binary data. Additionally, we have shown the importance of centering and that kernel centering can be used for the binary case.

### 3.4 Application to pharmacogenomics data

We applied the RV coefficient with both extensions to a collection of pharmacogenomics data (a combination of GDSC1000 (4) and MCLP (6), see Methods and Materials) to infer how the different datasets in this collection are related to each other. This collection consists of 3 binary datasets (mutation, CNA and cancer type) and 4 continuous datasets (methylation, gene expression, proteomics and drug response). Intersecting the objects that are present in all datasets resulted in data for 206 objects.

We used the PC algorithm (2; 12) (Supplementary Materials) to study the relationships between datasets. Briefly put, this algorithm starts out with a fully connected graph, where each node corresponds to a dataset, and removes the edge between two datasets **X**_1_ and **X**_2_ when *RV*(**X**_1_, **X**_2_|C) ≈ 0 (i.e. when it is not significantly different from 0). This step is repeated for increasingly larger sets of C, from C = θ (no datasets) to C = U \ {**X**_1_,**X**_2_} (all datasets except **X**_1_ and **X**_2_), until either the edge is removed or all possible sets have been assessed. Finally, the PC algorithm attempts to, under certain assumptions, determine the directionality of the edges. However, for the pharmacogenomics data, the algorithm was unable to infer the directionality of any edge in the graph.

Using the approach outlined above, the PC algorithm essentially summarizes the set of all 560 partial matrix correlations in a topology. An important caveat of this approach is that it uses the absence of a significant association to determine the absence of a relation between two datasets. As this may not always be true (there may be such a relation, but we may not have enough objects to detect it), we will also inspect the underlying (partial) matrix correlations and their confidence intervals for the most important hypotheses generated from the topology.

**Fig. 5.**
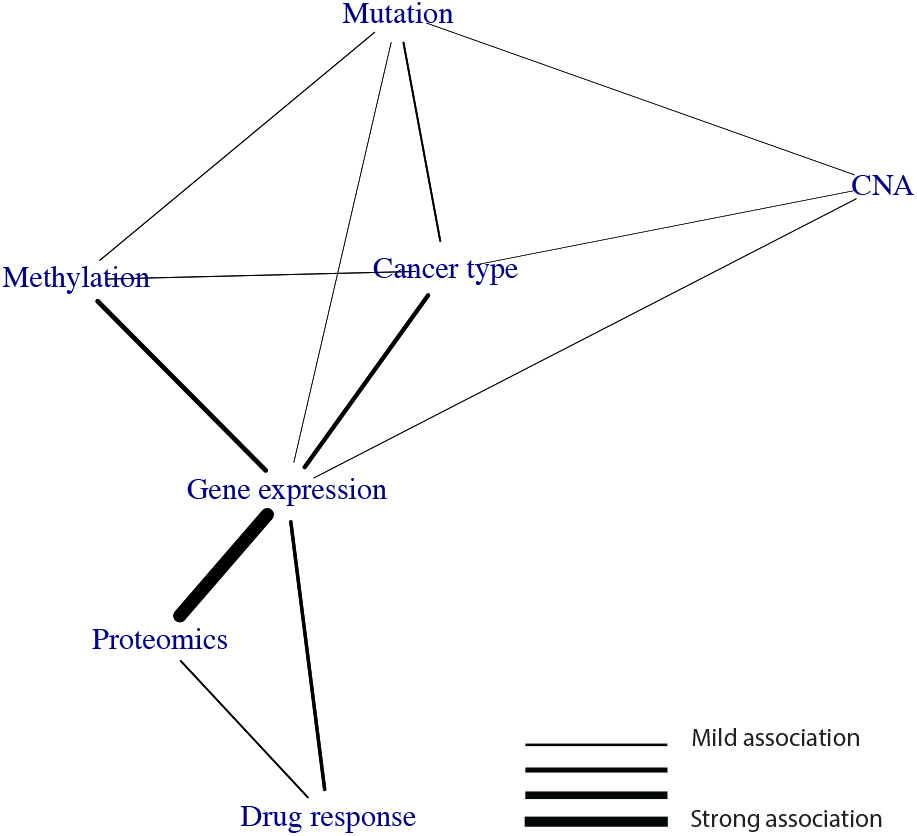
Relationships between datasets in the pharmacogenomics data, as determined using the PC algorithm run on the partial matrix correlations. An edge indicates that two datasets share information that is not present in any of the other datasets.

**Fig. 6.**
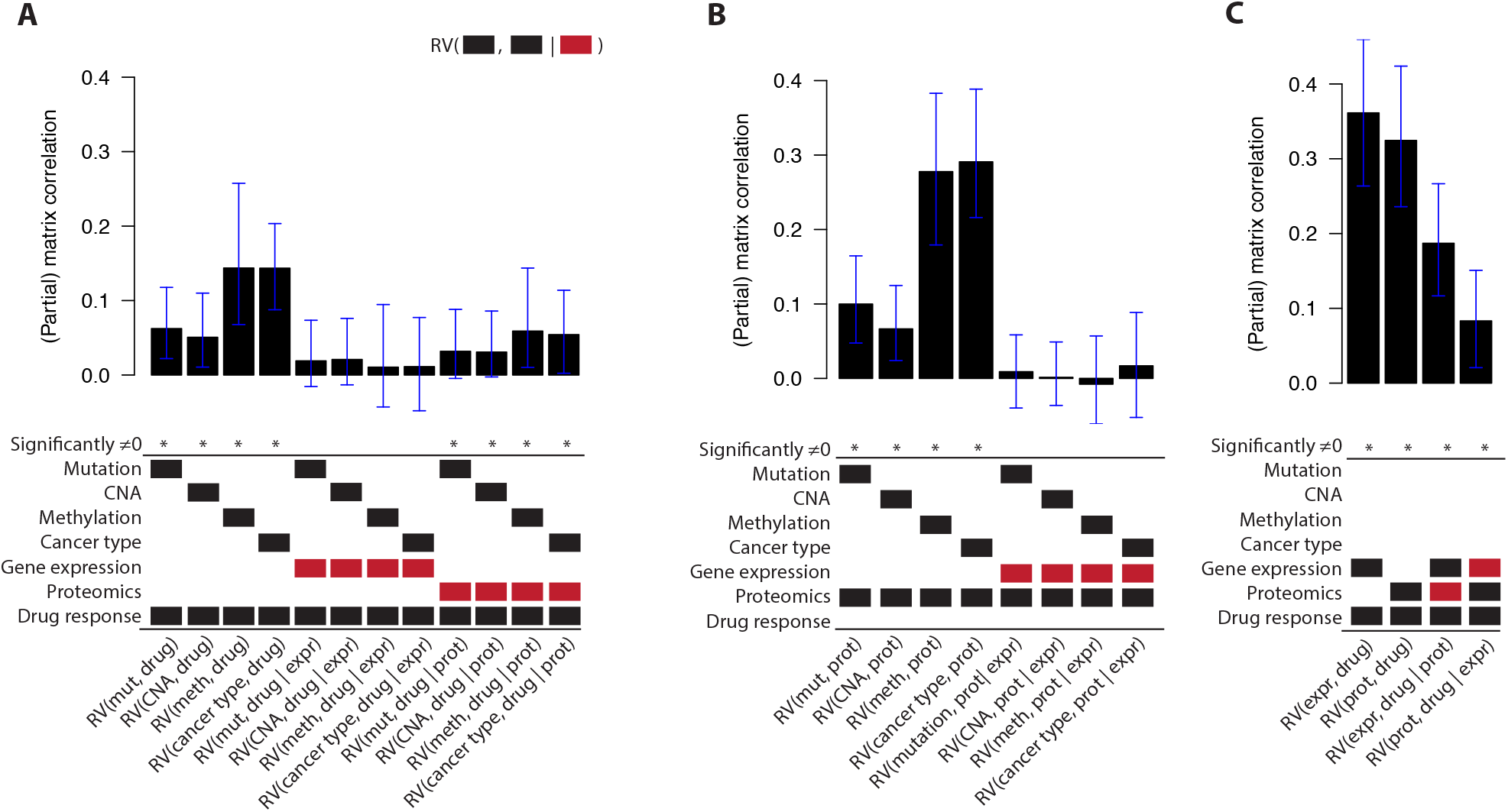
The (partial) matrix correlations for different *RV*(X_1_, X_2_|X_3_) in the pharmacogenomics data. For each bar in the barplot, X_1_ and X_2_ are indicated by the black blocks, and X_3_ is indicated by the red block. A (partial) matrix correlation was significant when p < 0.01. The error bars indicate the 99% confidence interval. Abbreviations: mut, mutation; meth, methylation; expr, gene expression; prot, proteomics; drug, drug response.

Figure 5 shows the topology resulting from the PC algorithm. Gene expression takes up a strikingly central position in the graph, being connected to all other data types. Using the underlying partial correlations and their confidence intervals, we verify that gene expression acts as a mediator between the ‘upstream data’ (mutation, CNA, methylation and cancer type) on the one hand and the drug response data on the other hand: the partial matrix correlations between these datasets and the drug response drop to nearly zero when correcting for gene expression (Figure 6A).

Proteomics also takes up an interesting position in the graph. The proteomics data shows a very strong relationship with gene expression (*RV* = 0.76). Interestingly, using the underlying partial matrix correlations, we see that this relationship fully contains the information shared between the upstream data and proteomics: *RV* (**X**_*i*_, proteomics | expression) ≈ 0, for each dataset **X**_*i*_ in the upstream datasets (Figure 6B). Finally, gene expression and proteomics share information with drug response that is not present in the other dataset: *RV* (expression, drug response | proteomics) > 0 and *RV* (proteomics, drug response | expression) > 0 (Figure 6C). Hence, even though gene expression and proteomics share a large amount of information, they both contain unique information with respect to drug response.

Overall, we have shown here that our methodology can be used to infer how different datasets are related to each other.

### 3.5 Identifying which variables predictive of drug response are distinct to either gene expression or proteomics

The topology that we have inferred suggests that for accurate prediction of drug response we only need gene expression and proteomics. Indeed, when we train Elastic Net models (17) (Supplementary Materials) to predict the drug response from either all datasets (other than drug response) or from only gene expression and proteomics, we found that they result in virtually identical predictive performance (Supplementary Figure 2A).

We then asked which variables are both predictive of drug response and distinct to either gene expression or proteomics. To answer this question, we used TANDEM (1) (Supplementary Materials). Briefly, given a response vector *y* (e.g. drug response of a single drug) and two datasets **X**_1_ and **X**_2_ (e.g. gene expression and proteomics), TANDEM uses two stages of Elastic Net regression to first identify all variables in **X**_1_ that are associated with y, and then identify all variables in **X**_2_ that are associated with y but whose information is not present in **X**_1_.

For each drug, we trained two TANDEM models:

- GEX_unique_: a model that uses proteomics in the first stage and gene expression in the second stage, thereby identifying variables with information that is unique to the gene expression data.
- PROT_unique_: the counterpart of GEX_unique_, with gene expression in the first stage and proteomics in the second stage.

We found that GEX_unique_ mostly uses proteomics data and PROT_unique_ mostly uses gene expression data, while both achieve similar predictive performance (Supplementary Figure 2B-D. This is of course not very surprising, as we have already seen using the RV coefficient that a lot of information is shared between the gene expression and proteomics data.

For each drug and for both TANDEM models, we then determined variable importance scores (Supplementary Materials) and averaged these over drugs to identify variables that made the largest overall contribution to the prediction of drug response. For GEX_unique_, the most important gene expression variable was ABCB1 expression. ABCB1 is a protein in the cell membrane that pumps foreign substances (including drugs) out of the cell. As such, it is known to be associated with resistance to a wide range of drugs (3). The proteomics data we considered here did not contain ABCB1, hence it is not unexpected that this information is not present in the proteomics data.

For PROT_unique_, the most important variable was MEK1 S217/S221 phosphorylation (pMEK1). The phosphorylation of MEK1 indicates MAPK pathway activation and is hence associated to sensitivity to MAPK pathway inhibitors, such as BRAF, MEK and ERK inhibitors. As the proteomics data contains both phosphorylation and protein abundance variables, we wondered whether one of these classes might be enriched in the distinct proteomics – drug response part. However, we found no significant difference between the variable importance scores in the PROT_unique_ models for these two classes (*p* = 0.68, Mann-Whitney U Test) (Supplementary Figure 2E).

Altogether, we have shown here that, informed by the topology of the datasets we inferred with iTOP, we can identify which variables correspond to distinct gene expression– drug response and proteomics–drug response relationships.

## 4 Discussion

In this work, we have introduced iTOP, a methodology to infer a topology of relationships between datasets. To this end, we have extended the RV coefficient for partial matrix correlations, allowing one to identify how much information is shared between two datasets, but not present in other datasets. In addition, we have also extended the partial RV coefficient for binary data, using the Jaccard coefficient. We have tested both extensions using artificial data and used them to infer a topology of the pharmacogenomics data. Finally, we have zoomed in on part of the topology and have identified variables predictive of drug response that are distinct to either gene expression or proteomics using TANDEM.

An important caveat of the PC algorithm used in our approach is that the absence of a significant p-value does not necessarily mean the absence of a relationship between two datasets: it can also mean this relationship is present, but that we did not have enough power to detect it. Of note, this also means that the inferred topology can change as the number of objects increases, simply because this enhances our ability to detect very small effects. For these reasons, we suggest to not solely rely on p-values to determine the absence or presence of these links. Instead, we suggest using the PC algorithm as a tool to summarize the results from the numerous possible partial matrix correlations into a topology, after which the hypotheses generated from this topology should also be assessed by inspecting the relevant (partial) matrix correlations and their confidence intervals. These values will give an indication of both the strength of the associations and how well we can estimate these, and may hence suggest the inclusion of an association that is strong but uncertain, or the exclusion of a certain – but weak – association.

We note that there are other options for binary similarity measures besides the Jaccard coefficient. For example, we have considered the phi coefficient, which is the Pearson correlation applied to binary measurements (15; 16). The main benefit of the phi coefficient is that it is a centered measure and hence kernel centering of the resulting configuration is not required. A minor disadvantage of the phi coefficient is that it is not defined in cases where objects consist of only zeroes or only ones. This can be easily circumvented however, for example by defining phi(x,y) = 0 in these cases. The main disadvantage of the phi coefficient lies in its definition of similarity: for the phi coefficient, both coinciding zeroes and ones contribute towards similarity, whereas for the Jaccard similarity only coinciding ones do. We believe objects are similar when they share the same mutations (rather than the absence of mutations) and hence prefer the Jaccard similarity here.

In future work, the RV coefficient could be further extended for other types of data. For example, a matrix with ordinal data could be converted into a configuration matrix using the Spearman rank correlation or the 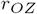 coefficient similarity (14; 16). Additionally, other semi-positive definite kernels that describe the similarity between objects could be used as a configuration matrix. For example, if we were to consider a dataset that is represented as a graph (where each node corresponds to an object), then a configuration matrix could be constructed using a graph diffusion kernel (5). Finally, as many multi-omics data contain patient survival data, defining a configuration matrix for survival data opens up interesting avenues for future research. For each of these extensions, careful assessment of the need of kernel centering will be required.

We believe that iTOP can be applied to a broad range of data, beyond the pharmacogenomics data analyzed here. Essentially, for all data in which the same objects have been characterized in multiple modalities, this methodology can be used to infer a topology of relationships between the resulting datasets. Hence, as multi-omics and phenotypic data is collected for increasingly more experiments, we believe our methodology will be highly relevant and widely applicable.

## Funding

The research leading to these results has received funding from the European Research Council under the European Union’s Seventh Framework Programme (FP7/2007-2013) / ERC synergy grant agreement nº 319661 COMBATCANCER.

